# Mutational signatures in wild type *Escherichia coli* strains reveal dominance of DNA polymerase errors

**DOI:** 10.1101/2023.03.02.530848

**Authors:** Sofya K. Garushyants, Mrudula Sane, Maria V. Selifanova, Deepa Agashe, Georgii A. Bazykin, Mikhail S. Gelfand

**Author notes:** Correspondence: Sofya K. Garushyants; Mikhail S. Gelfand.

## Abstract

While mutational processes operating in the *Escherichia coli* genome have been revealed by multiple laboratory experiments, the contribution of these processes to accumulation of bacterial polymorphism and evolution in natural environments is unknown. To address this question, we reconstruct signatures of distinct mutational processes from experimental data on *E. coli* hypermutators, and ask how these processes contribute to differences between naturally occurring *E. coli* strains. We show that both mutations accumulated in the course of evolution of wild type strains in nature and in the lab-grown non-mutator laboratory strains are explained predominantly by the low fidelity of DNA polymerases II and III. By contrast, contributions specific to disruption of DNA repair systems cannot be detected, suggesting that temporary accelerations of mutagenesis associated with such disruptions are unimportant for within-species evolution. These observations demonstrate that accumulation of diversity in bacterial strains in nature is predominantly associated with errors of DNA polymerases.

## Introduction

Mutagenesis provides the raw material for evolution, and understanding the processes involved in it is essential for studying the evolution of bacterial genomes. The progress in this area followed the rise of genome sequencing techniques, which allowed for genome-scale comparative analyses not restricted to reporter genes ^1–6^. Mutations occur in all cells at each replication cycle. However, mutation rates may differ even between laboratory strains of the same bacterial species^1^. Spontaneous mutations occur due to errors of enzymes involved in DNA replication or repair, combined with exogenous factors^7–9^. In bacteria, disruption of these processes yields hypermutator phenotypes, which are characterized by elevated mutation rates and changes in the mutational spectra^1,3,10^. Hypermutators accumulate point mutations at rates that are orders of magnitude higher than those of wildtype strains^11^.

Multiple distinct hypermutator phenotypes are known. The MutS, MutL, and MutH proteins contribute to hypermutator phenotypes as parts of the DNA mismatch repair system (MMR)^4^. In wildtype laboratory strains, mutations are substantially underrepresented in transcribed regions; however, this bias disappears in MMR-defective strains, which means that MMR preferably repairs transcribed regions^1,3^. A different type of hypermutators arise due to mutations in DnaQ, DNA polymerase III ε subunit that functions as a 5’-exonuclease ensuring the accuracy of chromosome replication^12,13^. On a rich medium, mutants that carry an almost inactive DnaQ have a 10 to 1000-fold higher mutation rate than the wildtype strains^14^. Inactivation of DnaQ also leads to activation of the SOS response^15^; however, SOS-induced error-prone polymerases do not contribute to the mutation rate or spectra of DnaQ mutants^5^, indicating that the changes observed in DnaQ mutants are caused by DnaQ inactivation itself. Furthermore, a mutator phenotype can arise from mutations in error-prone DNA polymerases IV (UmuDC) and V (DinB), and in DNA polymerase II (PolB). Inactivation of these polymerases yields elevated mutation rates under some conditions^16–18^ but not in normally growing cells^3^. Finally, inefficient removal of oxidized nucleotides, especially 8-oxoG, also influences mutation accumulation rates and spectra^3^. Oxidized guanines are removed via two major pathways, MutT that hydrolyzes 8- oxoG, or MutM or MutY that correct 8-oxoG:A mispairs^19,20^.

Previous studies of mutations in bacteria mainly considered single-nucleotide substitutions regardless of the adjacent nucleotides. Meanwhile, accounting for these nucleotides, the so-called mutational contexts, provides a finer resolution and allows to distinguish between processes that underlie mutagenesis. This approach has been widely applied and proved to be fruitful in cancer genomics^9,21–25^. Typically, it is assumed that individual processes affecting mutagenesis are characterized by specific patterns of mutations, or mutational signatures. A mutational signature as used here is characterized by the relative frequencies of six substitution subtypes: C>A (that is, the C:G pair to the A:T pair; hence equivalent to G>T), C>G, C>T, T>A, T>C, and T>G (equivalent, respectively, to G>C, G>A, A>T, A>G, and A>C) for each of the four possible adjacent 5’ and 3’ bases. This yields a total of 96 (6×4×4) possible mutation types. In cancer genomics, relative contributions of various mutational processes are inferred from tumor samples by decomposition of the observed mutational spectra into several standard signatures^20^. By contrast, in bacteria, mutational signatures may be observed directly by analyzing genomes of spontaneous mutator strains or laboratory knockouts of particular genes yielding mutation accumulation (MA) lines. Sequencing data on MA lines in model bacterial species are now available for strains generated in several laboratories in different set-ups, mostly for *Escherichia coli*^1,3,5,26–29^. Furthermore, data on naturally occurring mutators can be obtained from *in vitro* evolution experiments. For example, in the *E. coli* Long-term Evolution Experiment (LTEE)^4,30^, six strains acquired spontaneous mutations that affect MMR or removal of oxidized nucleotides. Both MA lines and evolution experiments allow for observing thousands of single nucleotide substitutions in mutator lineages, providing sufficient resolution to characterize mutational signatures associated with various mutational mechanisms.

In hypermutator strains, evolution primarily reflects mutational biases, with minimal contribution of selection^30^, so that mutational processes may be directly inferred from whole-genome mutational patterns. By contrast, slower accumulation of mutations in non-mutator strains is shaped by selection, both positive and negative^30^, with most positions in bacterial genomes subject to negative selection. Still, intergenic regions evolve in an almost neutral regime^31,32^, or are at least less affected by negative selection^33,34^. This allowed us to use intergenic regions to infer mutational processes in non-mutator strains.

Here, we address two questions: whether the mutational signatures and mutational processes underlying them may be inferred from experimental data on hypermutators; and how these mutational processes contribute to accumulation of mutations in *E. coli* strains in laboratory and in nature.

## Results

### Meta-analysis of mutation accumulation lines in *E. coli*

To study mutations accumulated via different mutational processes, we collected all available data on mutation accumulation in mutator strains of *E. coli* (Materials and Methods). We used sequencing data from four main sources: (i) six mutator strains that had emerged in the LTEE experiment: Ara-1 and Ara+6 with the MutT deficiency and Ara-2, Ara-3, Ara-4 and Ara+3 with mutations in MMR- associated genes^4^; (ii) MA strains for all major mutational processes from the Foster laboratory^1,3,5^, (iii) mutY deficient MA strains^6^; and (iii) a set of MA strains deficient in MMR components MutH, MutL, and MutS obtained specifically for this study (Supplementary Figure 1; Materials and Methods).

The resulting dataset included data on multiple distinct mutational processes, and for many of these processes, data came from several sources. Specifically, strains with defective MMR systems were collected from three independent labs, and in total yielded almost 12000 mutations (Supplementary Figure 1). The MutT and MutY mutator strains came each from two independent labs and had respectively nearly 9000 and 5000 mutations. We also considered strains with defective genes encoding the epsilon subunit of DNA polymerase III (*dnaQ*), DNA polymerase II (*polB*), DNA polymerase IV (*umuDC*), nucleotide excision repair ATPase (*uvrA*), base excision repair (BER) endonucleases III and VIII (*nth* and *nei*, respectively), and exonucleases III and IV (*xthA* and *nfo*, respectively)^3,5^. In total, our dataset included 40694 mutations from 17 experiments (Supplementary Figure 1).

To describe the overall mutation types characteristic of mutational processes in our data, we first extracted single-nucleotide substitutions and combined complementary mutations, hence retaining six types of substitutions. MA lines with similar deficiencies, such as MMR, MutT, and MutY deficient strains obtained in different laboratories, had similar single-nucleotide mutational profiles, which means that the observed mutational spectra were independent of experimental conditions (Supplementary Figure 1). As shown earlier, MMR and DnaQ mutants were associated with transitions (C>T and T>C)^3^; MutY mutants generated mostly C>A transversions; whereas MutT mutants almost exclusively carried T>G transversions. Strains deficient in the *nth* and *nei* genes carried prevalent C>T substitutions. MA strains deficient in *polB, umuDC, uvrA, xthA*, and *nfo* had relatively flat mutational profiles with a higher prevalence of all six types of substitutions relative to the “wild type” laboratory *E. coli* strain (Supplementary Figure 1).

### Reconstruction of mutational signatures from hypermutators

Analyzing only substitutions without accounting for their contexts does not produce sufficient resolution to distinguish, for example, between the excess of C>T mutations caused by MMR and by errors of polymerase III. To distinguish between mutations caused by different mutational processes, we considered three-nucleotide contexts, again collapsing complementary mutations, for a total of 96 types of mutations. We reconstructed the mutational profiles for each strain independently (Supplementary Figure 2).

We compared the mutational profiles of each strain using cosine similarity (see Materials and Methods), and observed that they form distinct clusters (Fig. 1a). We divided all strains into seven groups based on similarity of their mutational profiles: MMR-deficient strains from the LTEE experiment; DnaQ MA strains; MMR-deficient MA strains; MA strains with defective MutY, MutT, BER endonucleases III and VIII; and PolB-deficient and similar MA strains. The clustering matched the type of deficiency rather than the source of data. Indeed, while the spectra of MMR-deficient strains from LTEE slightly differed from those of MA strains which had accumulated mutations over a short period of time, the profiles in these two groups of MMR-deficient strains were still highly similar.

**Figure 1.**
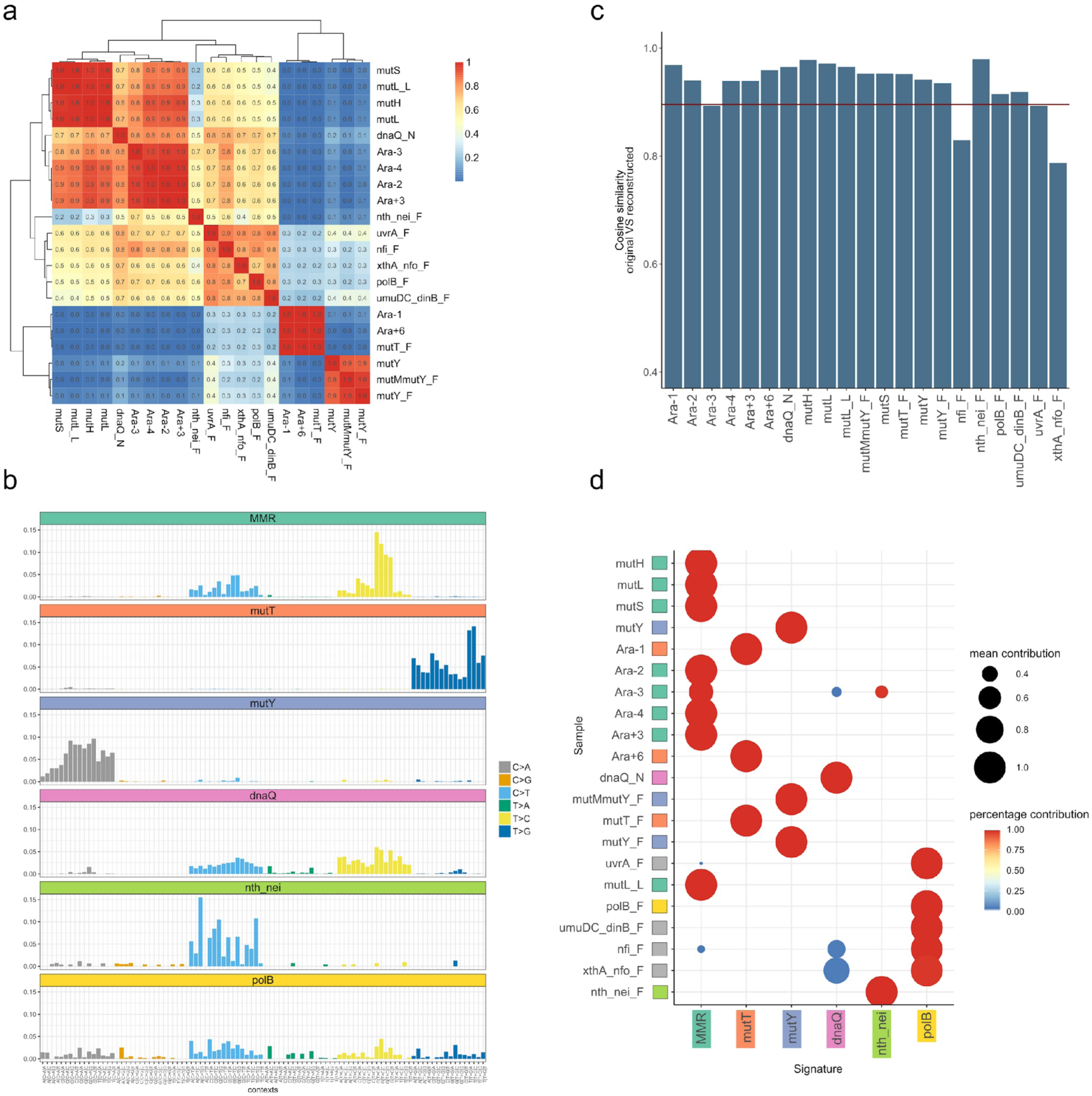
Reconstruction and validation of mutational signatures from the mutator data. (a) Cosine similarity matrix between 3-letter mutational profiles for various mutator strains. (b) Signatures of mutational processes reconstructed from the mutator strains (see also Supplementary Figure 4). (c) Similarity between the original and reconstructed profiles for the mutator strains. (d) Signatures of mutational processes are correctly assigned to the mutator strains. The size of the circle represents the mean contribution of the mutational signature in the sample, and the color of the circle represents the percentage of bootstrap replicates where this signature was observed.

To reconstruct the signatures characteristic of these mutational processes, we applied three different approaches. The direct approach was to define the mutational signature as the average mutational profile for all lines (both MA and LTEE) with a given defect. Additionally, we used two different techniques for *de novo* identification of signatures from a joint dataset of all lineages: non-negative matrix factorization (NMF) and Bayesian NMF (Materials and Methods). All three methods produced similar results (Fig 1b, Supplementary Figure 3, Supplementary Figure 4). The consistency between the direct and the *de novo* approaches provides, to our knowledge, the first direct experimental validation for the *de novo* approach. For further analysis, we used the signatures obtained directly from experimental data, assuming them to be the most reliable.

To ask how adequately the obtained signatures describe mutations observed in the mutators, we decomposed mutational profiles for each hypermutator strain into the signatures and then reconstituted the profiles. For 18 out of 21 lineages, the reconstructed profile had >90% cosine similarity with the original profile, and in the remaining cases, the similarity exceeded 80% (Fig 1c). The main process was correctly identified in all experimental setups (Fig 1d).

### Accumulation of mutations in non-mutator laboratory strains

To obtain mutational profiles for the strains accumulating mutations at a normal rate in laboratory experiments, we considered non-mutator phenotypes from the LTEE (two strains) and Foster (three strains) experiments (Materials and Methods). In total, these experiments comprised 1877 mutations across the genome, of which 40% were C>T transitions (Supplementary Figure 5a; Supplementary Table 1). The observed spectra differed from all profiles observed in the mutator strains, both for the whole genome and when we only considered intergenic regions (Supplementary Figure 5a,b). Accumulation of mutations in both LTEE mutators and MA lines was shown to be mostly driven by mutational bias and not affected by selection^35^, while in non-mutator strains, the whole genome mutational profile differed substantially from that of intergenic regions; specifically, it comprised considerably fewer T>C transitions (Supplementary Figure 5; Supplementary Table 1). The difference between the mutational profiles of coding and intergenic regions arises due to stronger selection and/or the activity of MMR in the coding regions. To control for these differences in our comparisons with mutator strains, for further analysis of non-mutators, we considered only 375 mutations observed in intergenic regions. This means that we were unable to study the contribution of MMR to mutation accumulation in non-mutator strains.

### Accumulation of mutations during the divergence of natural *E. coli* lineages

Finally, we considered mutational processes affecting natural populations of *E. coli*. For this, we collected a dataset of 522 complete *E. coli* genomes, reconstructed their phylogeny based on a set of single-copy universal genes, and selected four major clades each containing at least 40 genomes (see Materials and Methods), for a total of 470 genomes. These clades roughly corresponded to the established phylotypes A, B1, B2, and E (Figure 2; Supplementary Table 2). For each clade, we reconstructed the whole-genome alignment and inferred single-nucleotide substitutions at all phylogenetic branches with the baseml program of the PAML package ^36^. To minimize the effects of selection, we only considered substitutions in intergenic regions with more than 20000 substitution per phylogroup (Supplementary Table 1). To additionally confirm that the reconstructed mutational profiles are minimally affected by selection, we also separately analyzed intergenic regions between convergent genes (Supplementary Table 1), expected to be under weaker selection because they contain fewer regulatory sequences^33^. The reconstructed profiles were very similar between the considered clades, as were the profiles for all intergenic regions and intergenic regions of convergent genes (Supplementary Figure 6). In further analysis, we used the profiles built for all intergenic regions.

**Figure 2.**
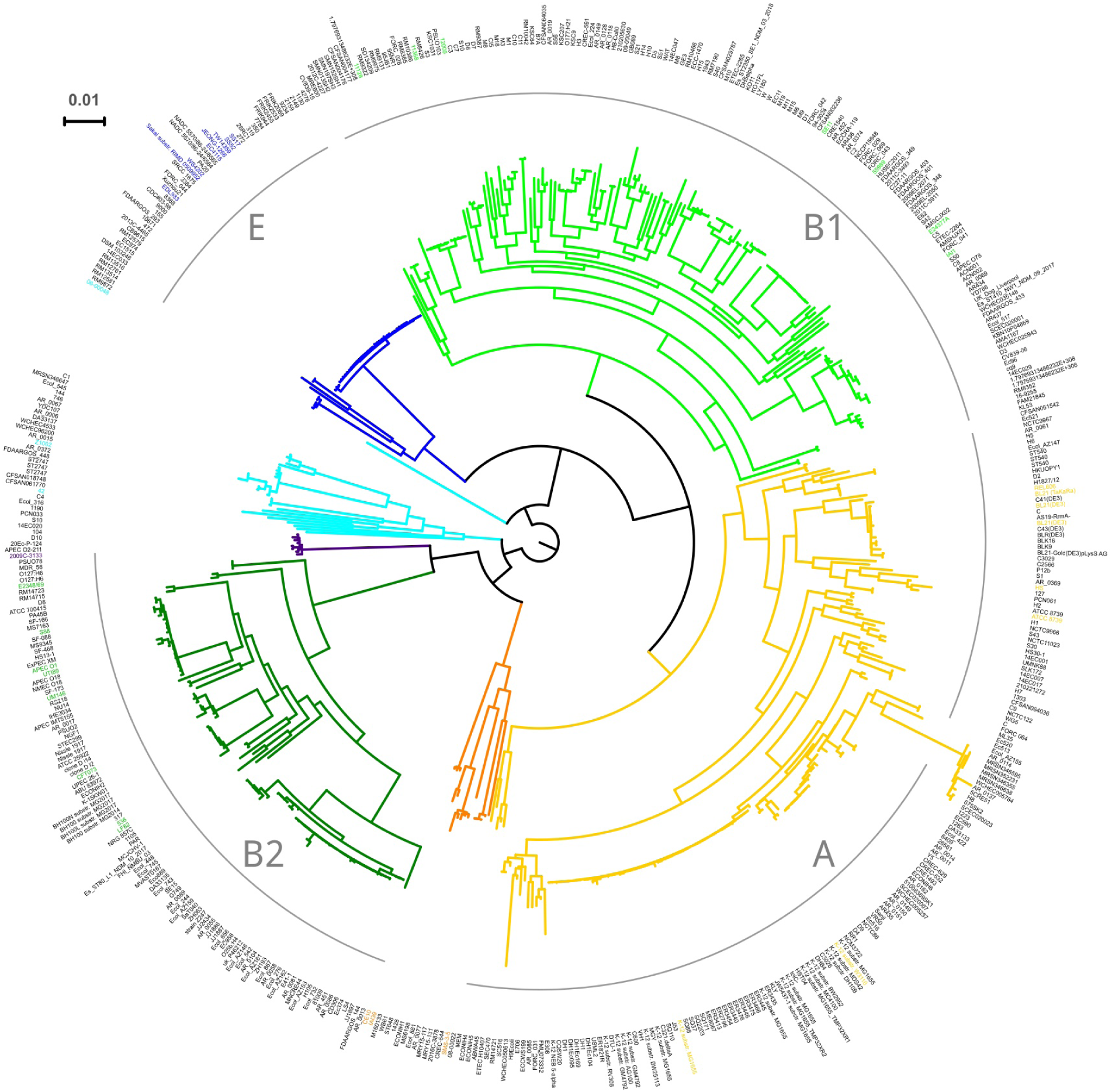
Phylogenetic tree of 522 complete *E. coli* genomes. The maximum likelihood tree was reconstructed from the concatenated alignment of universal single-copy orthologous genes (see Material and Methods). The four analyzed clades, A (yellow), B1 (green), B2 (dark green) and E (blue), are also indicated with grey arcs. Leaves for which the phylogroup is indicated in metadata are colored accordingly. Black leaves mean that no phylogroup information is available.

### Laboratory non-mutator strains and natural strains have similar mutational profiles

We find that the mutational profiles of intergenic regions of diverging natural *E. coli* lineages are very similar to those of lab-based non-mutator strains (Figure 3a). This indicates that accumulation of mutations in the course of evolution is driven by the same mutational processes as that observed in non-mutator strains during laboratory MA experiments. To better understand the processes responsible for mutation accumulation in these two settings, we decomposed the mutational profiles into a set of six signatures of six distinct mutational processes reconstructed from the mutator data.

**Figure 3.**
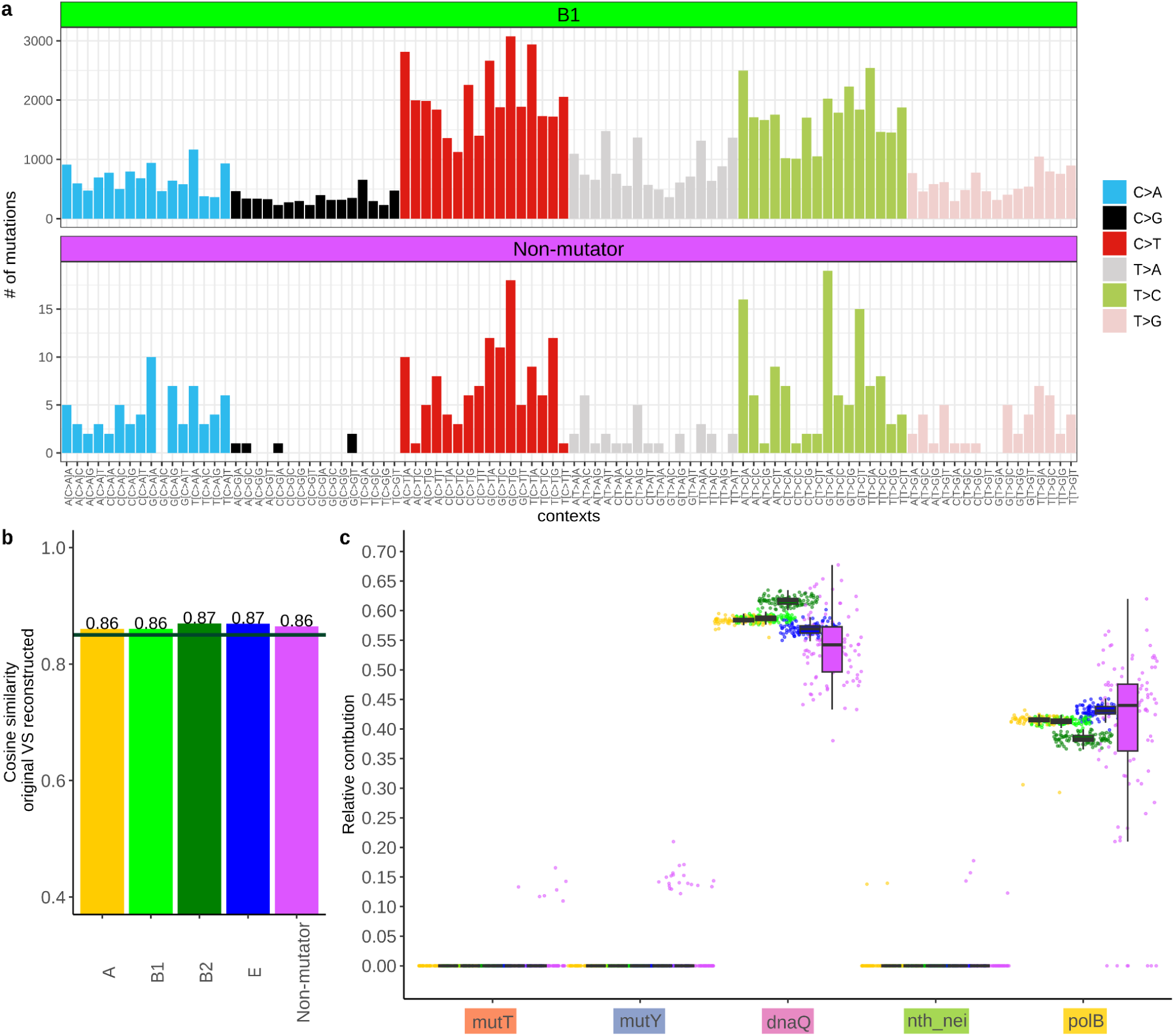
Contribution of mutational signatures to the evolution of intergenic regions in *E. coli* natural and in non-mutator laboratory strains. (a) Mutational profile for non-mutator strains and clade B1. (Profiles for all clades are shown in Supplementary Figure 6.) (b) Cosine similarity between the original mutational profiles and the reconstructed profiles. The horizontal line is at 0.85. (c) Relative contribution of each signature to the reconstructed profile. Each dot is the value obtained in one bootstrap replicate. Dots are colored by the data type as in (b).

Both the laboratory non-mutator strains and the natural lineages are rather well described by a combination of all six mutational processes, as indicated by the >86% cosine similarity between the actual mutational signature and its best reconstruction from these six processes (Figure 3b, Supplementary figure 7). The “missing” similarity in the natural lineages is very close to that in the laboratory settings and suggests that external and internal factors that are present in the natural environment of *E. coli* are present in laboratory conditions as well.

Consideration of contribution of each mutational process revealed that the predominant process causing mutations is associated with the DnaQ signature, which covered 59% (57%–61%) of all explained mutations in natural lineages and 56% (46%–65%) in laboratory non-mutators, suggesting a strong contribution of the major replicative DNA polymerase III. The remaining variability, comprising ∼45% of explained mutations, is explained by the PolB signature, indicating the contribution of the replicative DNA polymerase II. There was no signature of defective BER endonucleases III and VIII (Nth and Nei), MutT, or MutY (Figure 3c). As expected, we also saw zero contribution of the MMR mutational signature to both datasets (not shown in Figure 3d), as we considered non-coding regions, and MMR is only active in transcribed regions.

The inferred patterns were very similar between all four *E. coli* clades, indicating that they are characteristic of the overall *E. coli* evolution rather than lineage-specific peculiarities. The results were stable across bootstrap replicates, and robust when we considered only intergenic regions between convergent genes (Supplementary Figure 8). Additionally, we verified that the observed differences in the mutational profiles of natural strains and laboratory non-mutators were not an artifact of the ancestral state reconstruction by baseml. For that, counts for all identical mutations that had occurred at a particular position multiple times within a clade were set to one, e.g. recurrent identical substitutions at the same site were ignored. The resulting contributions of mutational signatures were almost identical to the ones described above for the full dataset (Supplementary Figure 9).

## Discussion

The properties of mutagenesis shape evolution, determining the load of deleterious mutations and channeling adaptation to new conditions. While the main processes resulting in single-nucleotide mutations are rather well understood, their relative contributions to bacterial evolution are largely unknown. In particular, it is not clear whether experimental laboratory conditions can adequately model mutagenesis in the complex natural environments experienced by bacterial lineages throughout their evolution. To address this question, we combine experimentally inferred data on mutagenesis with the mutational patterns revealed by lab MA experiments and with the reconstructed substitutions accumulated over thousands of years in evolving *E. coli* lineages.

This approach has several limitations. First, experimental studies of mutagenesis in bacteria are in their infancy. While tens of mutational signatures have been assigned to particular biochemical processes in higher eukaryotes such as humans^25^, the twelve mutational processes have been characterized in bacteria, but only six single-gene knockouts have sufficient data to be accurately described. The fact that the decomposition of the accumulated mutations into mutational signatures in our analysis (∼86% cosine similarity) is poorer than that in e.g. cancer genomics studies (>90%)^22^ may be partially explained by this deficiency. While our study reliably identifies the relative contributions of the studied mutational processes, knowledge of new mutational processes will further clarify the picture.

Second, inference of mutagenesis from MA lines and divergence can be confounded by selection. Bacteria have compact genomes, more than 80% of which are coding, meaning that the bulk of a genome is subject to selection; to a weaker degree, selection also affects intergenic regions^33,34^. While polymorphism and divergence data can be used to infer mutation patterns^37–39^, the effect of selection on the substitutions that distinguish long-term-evolved lineages is expected to be stronger than that in MA lines (where it is very weak^35^), as these mutations have survived within populations for a longer time and selection has had more opportunity to act. To mitigate this problem, we focused on genomic regions that are predicted to be least affected by selection. Most of our analyses were performed on intergenic regions, with the side effect that we were unable to study the contribution of the MMR system that mostly acts within transcribed regions. We also considered separately the regions between convergent genes, where the effect of selection was minimal^33^. The results were the same as for all intergenic regions, indicating that the effect of selection was low.

Third, our divergence analysis relies on correctness of the inferred phylogeny and the ancestral state. Systematic biases at these stages of the analysis could distort the inference of mutational spectra. However, the phylogenetic distances between the compared strains were relatively low (Fig. 2), and an alternative method of dealing with recurrent substitutions at a site yielded very similar results, indicating that our results were robust in this respect.

Given these limitations, we described the mutational signatures for six determinants of spontaneous mutagenesis in *E. coli*. Accounting for mutational contexts allowed us to quantitatively assess the causes of spontaneous mutations in the studied bacterial strains. Decomposing the spectra of mutations accumulated in the wild-type non-mutator strains in the laboratory, and in the natural environments over the course of evolution, into the signatures of six mutational processes allowed us to infer the prevalent forces responsible for mutagenesis.

We find that the mutational spectra of reconstructed substitutions that have occurred in the course of evolution of natural *E. coli* lineages and in laboratory non-mutator *E. coli* MA strains were similar to each other. This observation argues against the notion that *E. coli* evolution in the natural environment involves alternating episodes of accelerated evolution in the mutator regime and slow evolution in the non-mutator regime, suggesting instead that at the scale of evolution of different *E. coli* strains, the contribution of atypical mutational regimes to the overall mutation accumulation is minor. This is probably due to the high load of deleterious mutations associated with such a regime^35^. Long-term evolution at the level of bacterial species, which involves gains and losses of repair-associated genes, could follow a different pattern^6^.

The observed spectra are similar to those associated with disruption of DnaQ and PolB. DnaQ is a core epsilon subunit of DNA polymerase III that is required for proof-reading and is essential for accurate DNA replication in *E. coli*^40–42^. PolB is a supplementary DNA polymerase II; while it can be induced by the SOS response^43,44^, it is also thought to be involved in DNA repair and to play an auxiliary role in chromosomal DNA replication under normal conditions^45^. The fact that the mutational spectra characterizing real-life evolution largely coincides with those of the enzymes involved in replication- associated repair indicates that the accumulated mutations occur co-replicatively, and suggests that they are associated with insufficient efficiency of these two repair enzymes during normal replication.

The near-identical contributions of PolB and DnaQ signatures to both natural *E. coli* lineages and laboratory non-mutator *E. coli* MA strains supports the notion that the bulk of the mutations associated with these two signatures occur co-replicatively. Indeed, if one of these components had a substantial fraction of non-co-replicative mutations, we would expect a higher contribution of the corresponding enzyme to the mutation spectra of natural strains, and lower, to the laboratory strains. This is because laboratory strains have a much higher replication rate than natural strains (20 minutes vs. 15 hours per generation^46^), and the proportion of co-replicative vs. “clock-like” mutations in the former should be higher.

Surprisingly, no contribution of incorrect removal of 8-oxo-G, caused by ionizing radiation or oxidative metabolism, was observed in either setting, indicating that the time spent by bacteria in oxidized conditions is probably low.

In summary, while it has been disputed whether evolutionary experiments in monoculture bacteria adequately represent the real evolution in nature^47^, when the complexity of the real world is removed, it appears that such experiments in *E. coli* can very well represent mutation accumulation in natural environments. It would be interesting to expand this analysis to different evolutionary settings and other bacterial species.

## Materials and Methods

### Bacterial strains

We obtained the wild-type (WT) strain of *E. coli* K-12 MG1655 from the Coli Genetic Stock Centre (CGSC, Yale University), streaked it on LB (Luria Bertani) agar, and chose one colony at random as the WT ancestor for subsequent experiments. We similarly obtained the mutator strains of *E. coli* (ΔmutH, ΔmutL, ΔmutS) from the Keio collection (BW25113 strain background)^48^ of gene knockouts from the same stock centre. These gene knockouts were made by replacing open reading frames with a Kanamycin resistance cassette, such that removing the cassette generates an in-frame deletion of the gene. The design of gene deletion primers ensured that downstream genes were not disrupted due to polar effects. For each strain, we moved the knockout loci from the BW25113 background into the MG1655 (WT) background using P1-phage transduction^49^. We then removed the kanamycin resistance marker by transforming kanamycin-resistant transductants with pCP20, a plasmid carrying the flippase recombination gene and Amp^R^ resistance marker. We grew ampicillin resistant transformants at 42 ºC in LB broth overnight to cure pCP20. We streaked out 10 μL of these cultures on LB plates. After 24 hours, we replica-plated several colonies on both kanamycin-LB agar plates and ampicillin-LB agar plates, to screen for the loss of both kanamycin and ampicillin resistance. We PCR-sequenced the knockout locus to confirm removal of the kanamycin cassette.

### Experimental evolution under mutation accumulation (MA) and whole-genome sequencing to identify mutations

For each mutator strain, we founded 20 MA lines from a single ancestral colony (two lines per Petri plate), incubated at 37ºC, as described earlier for wild-type MA^50^. For each line, every 24 hours we streaked out a random colony (closest to a pre-marked spot) on a fresh LB agar plate. Every 4-5 days, we inoculated a part of the transferred colony in LB broth at 37ºC for 2-3 hours and froze 1 mL of the growing culture with 8% DMSO at -80 ºC. For the current study, we used stocks frozen on day 50 (∼1375 generations).

For each strain, we sequenced whole genomes from the last stored colony of each lineage as follows. We inoculated 2 μL of the frozen stock of each evolved MA line (or the ancestor) in 2 mL LB, and allowed the cells to grow overnight at 37ºC with shaking at 200 rpm. We extracted genomic DNA (GenElute Bacterial Genomic DNA kit, Sigma-Aldrich), quantified it (Qubit HS dsDNA assay, Invitrogen), and pooled equal amounts of genomic DNA from each of the 20 lines for a given mutator strain into a single tube. Thus, we prepared 3 pools of genomic DNA, one each for the mutators Δ*mutH*, Δ*mutL*, and Δ*mutS*, each containing an equal amount of genomic DNA from each of the 20 MA lines. We then prepared paired-end libraries from the Δ*mutH*, Δ*mutL* and Δ*mutS* MA ancestors and the three pooled genomic DNA samples using the Illumina Nextera XT DNA library preparation kit per the manufacturer’s instructions. We sequenced all libraries on the Illumina MiSeq platform using the 2×250bp paired-end V2 reaction chemistry. For the Δ*mutH*, Δ*mutL* and Δ*mutS* MA ancestors, we obtained 1.14M, 1.49 M and 1.06 paired end reads with quality > Q30 respectively. For the pooled evolved lines, we obtained 2.09M, 2.18M, and 2.35M paired-end reads respectively with quality > Q30, corresponding to an average per base coverage of ∼ 116x, 121x, and ∼130x. For each sample, we aligned quality-filtered reads to the NCBI reference *E. coli* K-12 MG1655 genome (RefSeq accession ID GCA_000005845.2) using the Burrows-Wheeler short-read alignment tool. We generated pileup files using SAMtools^51^ and used VARSCAN to extract a list of base-pair substitutions and short indels (<10bp)^52^. For the MA ancestors, we only retained mutations with >80% frequency that were represented by at least 5 reads on both strands for further analysis. For the pooled evolved MA lines, we retained mutations with > 5% frequency that were represented by at least 3 reads on both strands. After removing ancestral mutations (i.e. mutations that differentiated the ancestors from the reference *E. coli* genome) from the evolved lines, we identified 985, 875 and 768 SNPs in ΔmutH, ΔmutL and ΔmutS evolved MA lines respectively.

### Hypermutator data published earlier

We considered three main sources of data. (1) Sequences of six mutator strains from the LTEE experiment: Ara-1 and Ara+6 with MutT deficiency and Ara-2, Ara-3, Ara-4, and Ara+3 with mutations in MMR-associated genes^4^. The data for this experiment were downloaded from http://barricklab.org/shiny/LTEE-Ecoli/. Only SNPs were selected from the obtained file. Additionally, mutations observed at different time points for the same strain were merged to avoid repeated counting of mutations that had occurred early in the experiment. The obtained table was converted to the vcf format. (2) Experiments on MA strains for all major mutational processes from Patricia L. Foster’s laboratory^1,3,5^. In case of data from refs. ^1,3^ we obtained information about mutations from the Supplementary Materials and converted it to the vcf format. To obtain mutations caused by inactivated DnaQ^5^, we downloaded sequencing data from SRA (SRA IDs: SRR7748246, SRR7748247, SRR7748248, SRR7748249, SRR7748253, SRR7748306, SRR7748326, SRR7748342, SRR7748539, SRR7748540, SRR7748541, SRR7748542, SRR7748543, SRR7748592, SRR7748593, SRR7748594, SRR7748595, SRR7748733, SRR7748734, SRR7749018, SRR7749020, SRR7749022, SRR7749036, SRR7749037, SRR7749062, SRR7749108). The obtained reads were trimmed with Trimmomatic- 0.39^53^ with parameters LEADING:3 TRAILING:3 MINLEN:36. The trimmed paired reads were processed as described in the original publication. The variants were called with mpileup from bcf- tools with the minimal sequencing depth of at least 20 reads. In total, we obtained 13512 SNPs of which four occurred more than once and were discarded, yielding 13504 mutations.

### Laboratory strain data

We used non-mutator data from LTEE^4^, and experiments from Patricia L. Foster’s laboratory^3^. All LTEE non-mutator strains were merged together for further analysis. Duplicated mutations were filtered out to avoid selection effects. We considered separately all available mutations and mutations in intergenic regions.

### Reconstruction of the *E. coli* phylogeny

Complete genomes of 522 *E. coli* strains were downloaded from Genbank^54^. The complete list of genomes is provided in Supplementary Table 1. The initial set of orthology groups (OGs) from 32 phylogenetically diverse *E. coli* and *Shigella* strains was taken from^55^. We produced alignments for each universal group (OGs with single-copy genes present in all analyzed genomes) with ClustalW version 2.1^56^. For each alignment, we generated a consensus sequence with the EMBOSS package ver. 6.6.0 and mapped these consensuses to the genomes with bowtie2^57^. All groups that remained universal and single copy (690 genes) were realigned. Alignments for individual genes were concatenated and all columns with gaps were removed. This final alignment was used to construct the phylogenetic tree with PhyML v3.3 with the GTR substitution model^58^.

### Genome alignments and reconstruction of the ancestral states

Based on the *E. coli* phylogenetic tree, we selected four clades that contained at least 45 genomes. For the genomes where phylogroup information was available (shown as colored leaves in Figure 2), we checked that genomes within the clade belong to the same *E. coli* phylogroup.

For each of the selected clades, we constructed genome alignments with ProgressiveCactus^59^. For each clade, we selected one representative genome that was then used to identify gene boundaries and select intergenic regions between convergent genes (Supplementary Table 1). The output format was converted to maf, with the representative genome serving as the reference, using the hal2maf tool from ProgressiveCactus with parameters: --noDupes --onlySequenceNames --noAncestors --maxBlockLen 1000000. From this alignment, we extracted all variable columns in intergenic regions that (i) contained at least five genomes and (ii) had conserved columns to their left and right. Variable columns with adjacent variable columns or columns with gaps were discarded.

For these positions, we reconstructed the ancestral state with the baseml program from the PAML package v4.9j^36^.

### Reconstruction of mutational profiles in natural strains

From the PAML results (rst), we obtained the number of substitutions in the variable column for each site and then pooled this for all sites for each context within the clade.

We calculated the number of substitutions in all intergenic regions, and separately in intergenic regions between convergent genes. In order to make results compatible between different clades, the number of observed mutations in each context was normalized by the total number of mutations observed for the clade. To control for possible incorrect reconstruction of ancestral states by baseml, in an alternative procedure, if there were multiple identical (parallel) substitutions at the given site for the particular clade, we counted them as just one substitution in mutation counts.

### Mutational signatures

For each available experiment we calculated the contexts based on the initial strains in which the experiment was performed. Mutational signatures reconstructed from the data were calculated as the average normalized contribution of a particular substitution among all available mutator strains with the given deficiency. The reconstruction of mutational signatures was performed in R ver 4.1.1. To reconstruct signatures with the NMF approach, we used the NMF package in R. The Bayesian NMF approach was implemented with the ccfindR package. The scripts to reproduce this analysis are available on github (see Data Availability).

Deconvolution of obtained mutational profiles by mutational signatures was performed with the MutationalPatterns package^60^.

## Authors contributions

SKG and MSG conceived the project. MS performed the MA experiments, sequencing and variant calling from the data. SKG and MVS reconstructed mutational signatures from experimental data. SKG performed data analysis. DA, GAB and MSG supervised the research. SKG, GAB, and MSG wrote the manuscript with contributions from all authors.

## Acknowledgements

The project was initiated with Arina Kolotova at the Summer School of Molecular and Theoretical Biology for high school students. We thank our colleagues, in particular Dr. Anna Kaznadzey (IITP RAS) and Dr. Olga Vakhrusheva (Skoltech), for useful discussions and suggestions. We acknowledge help and support from the Next-generation genomics facility at the National Centre for Biological Sciences (NCBS) for whole-genome sequencing of bacterial strains. This work was supported by the Russian Science Foundation (18-14-00358), the Council for Scientific and Industrial Research (CSIR-India; senior research fellowship to MS), the National Centre for Biological Sciences (NCBS-TIFR), and the Department of Atomic Energy, Government of India (Project Identification No. RTI 4006).

## Data availability

All data and scripts required for the analysis of data are deposited in github: https://github.com/garushyants/Ecoli_MutationalSignatures

