## Supplementary_data for "Mutational signatures in wild type *Escherichia coli* strains reveal dominance of DNA polymerase errors"

[2] Present address: National Center for Biotechnology Information, National Library of Medicine, National Institutes of Health, Bethesda, MD, USA

Supplementary Table 1. Number of observed mutations in intergenic regions by substitution type in divergence and non-mutators.

Supplementary Table 2. *Escherichia coli* genomes used for divergence analysis

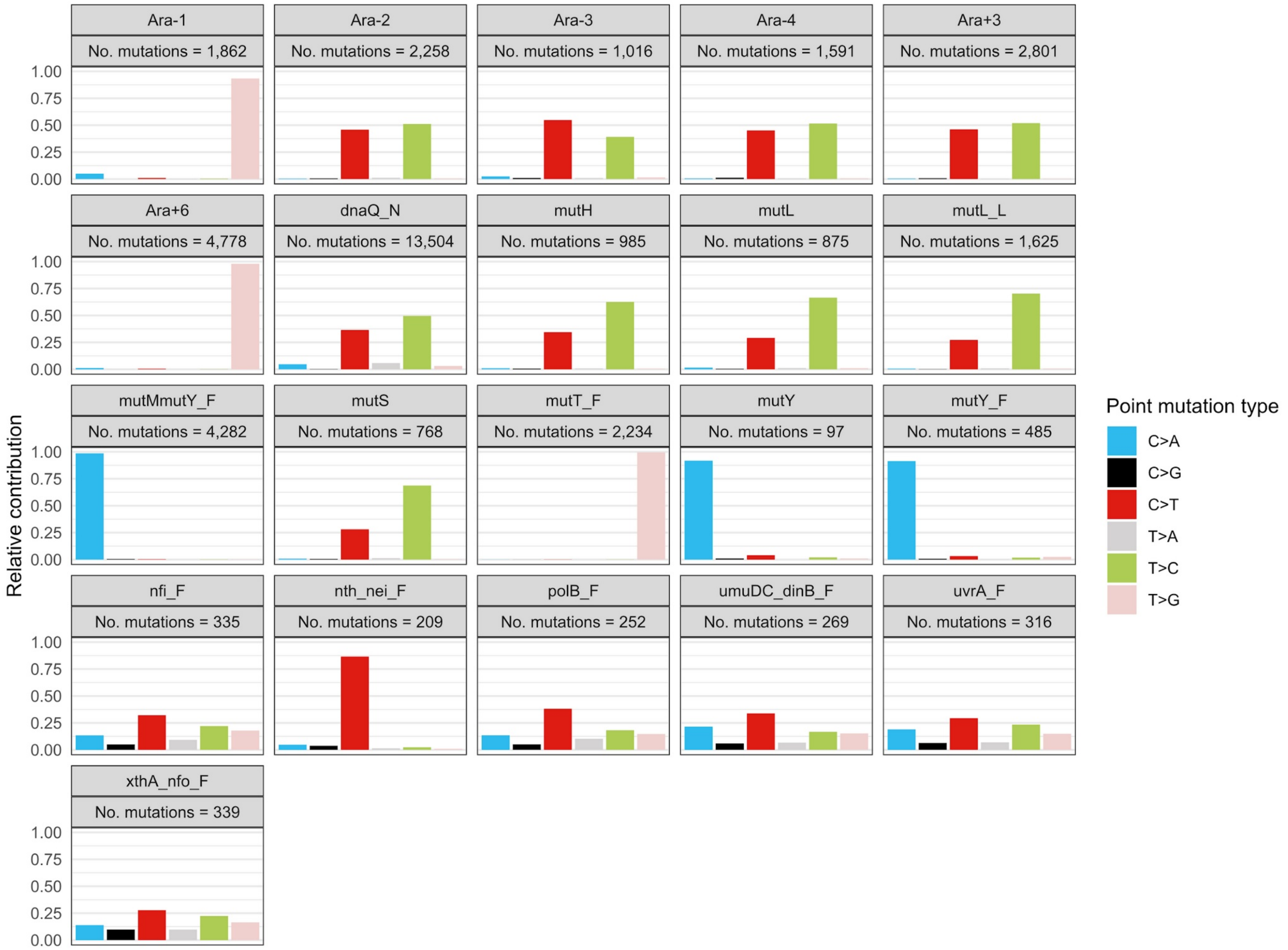


Supplementary Figure 1. Mutations observed in experimental data from various sources.

Ara-1 and Ara+6 with mutT deficiency and Ara-2, Ara-3, Ara-4 and Ara+3 with mutations in MMR associated genes (Tenaillon et al., 2016), the experiments on mutation accumulation strains for major mutational processes from Patricia L. Foster laboratory dnaQ_N. mutL_L,mutMmutY_F,mutT_F,mut_F,nfi_F,nth_nei_F,polB_F,umuDC_dinB_F, uvrA_F,xthA_nfo_F (Lee et al., 2012; Foster et al., 2015; Niccum et al., 2018), mutY from Sane et al, 2022, biorxiv; and also data on mutation accumulation strains with defective MMR genes *mutH*, *mutL* and *mutS and* obtained in this study.

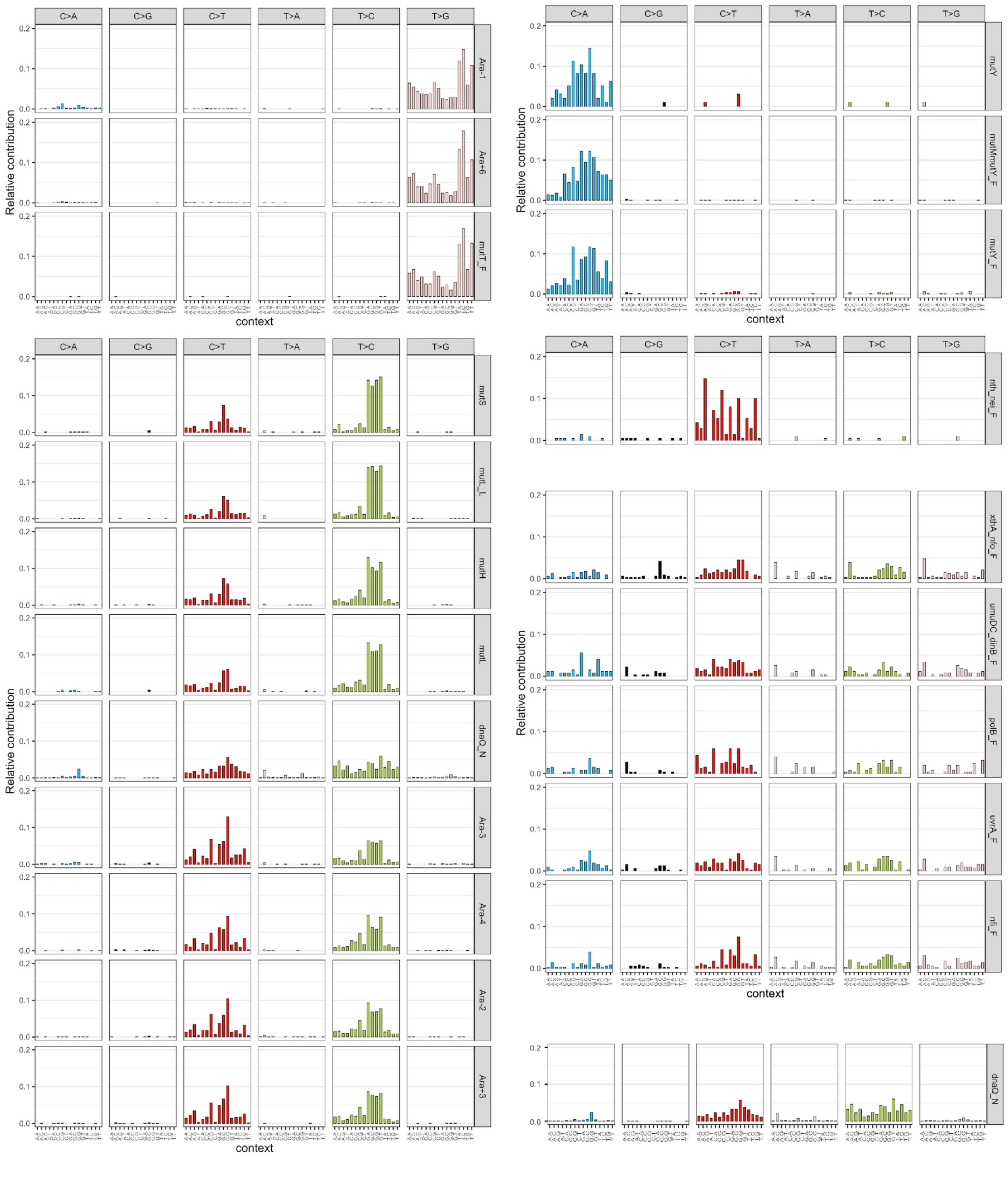


Supplementary Figure 2. Trinucleotide substitution profiles for various mutator strains grouped by profile similarity.

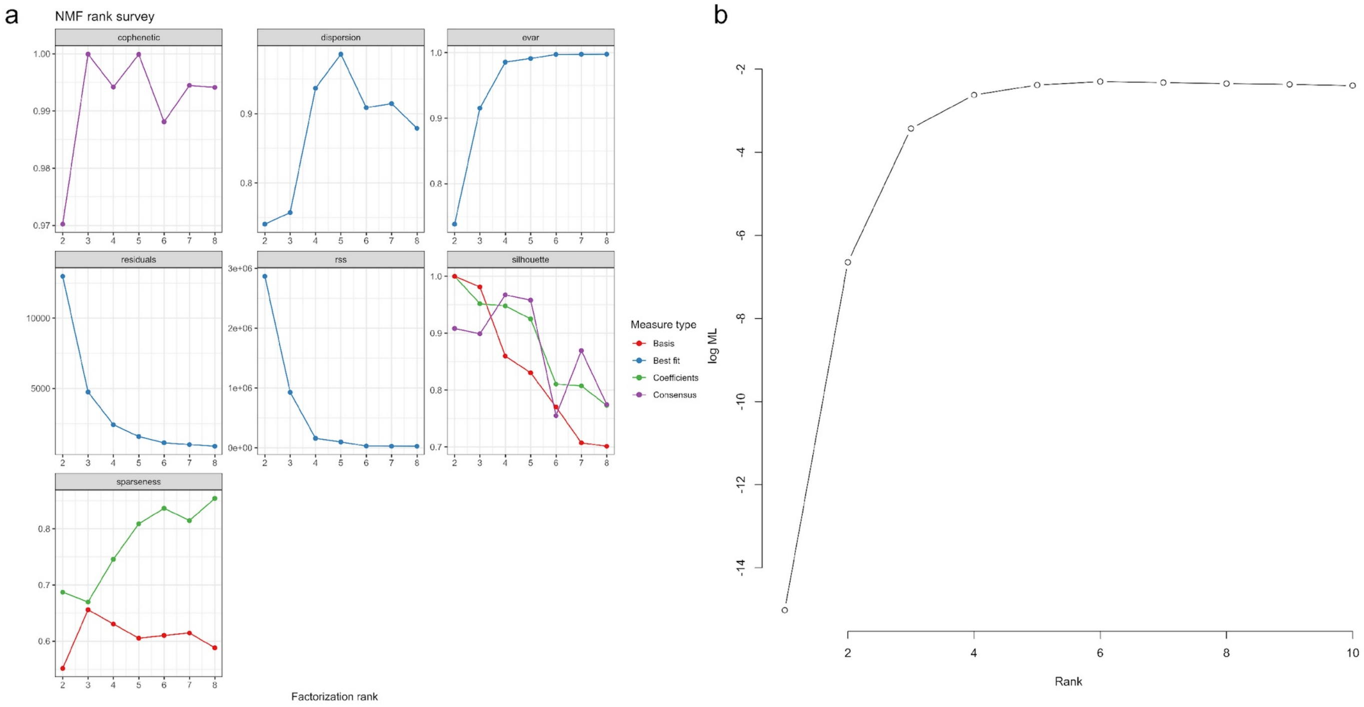


Supplementary Figure 3. Estimation of the number of mutational signatures in the data from hypermutator strains. (a) Estimations for negative matrix factorization (NMF) method. (b) Estimation for Bayesian NMF

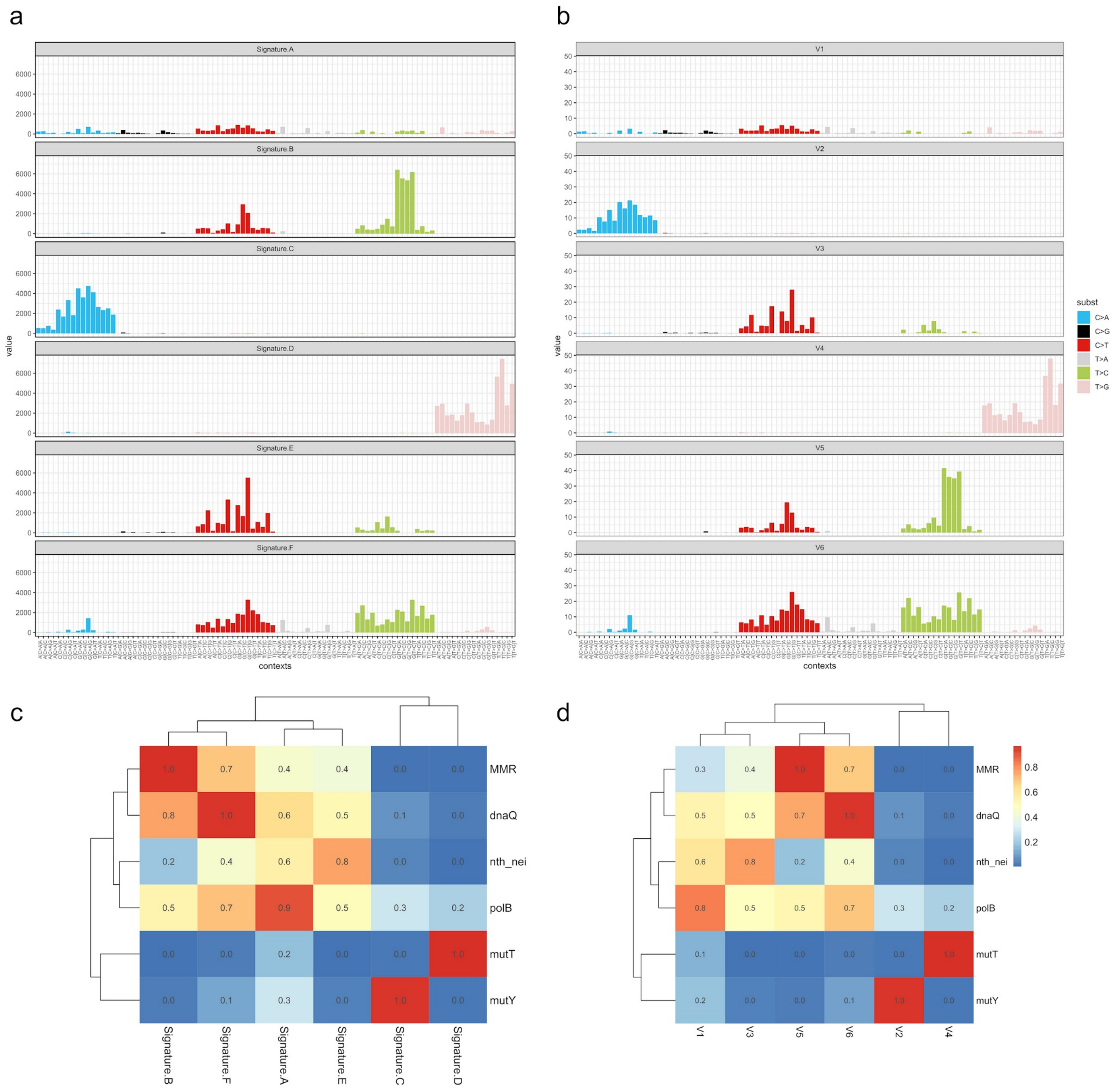


Supplementary Figure 4. Mutational signatures reconstructed by various methods (a) Profiles of signatures reconstructed by NMF (c) Profiles of signatures reconstructed by Bayesian NMF. (c) Comparison of signatures reconstructed by NMF and obtained from the data. (d) Comparison of signatures reconstructed by Bayesian NMF and obtained from data.

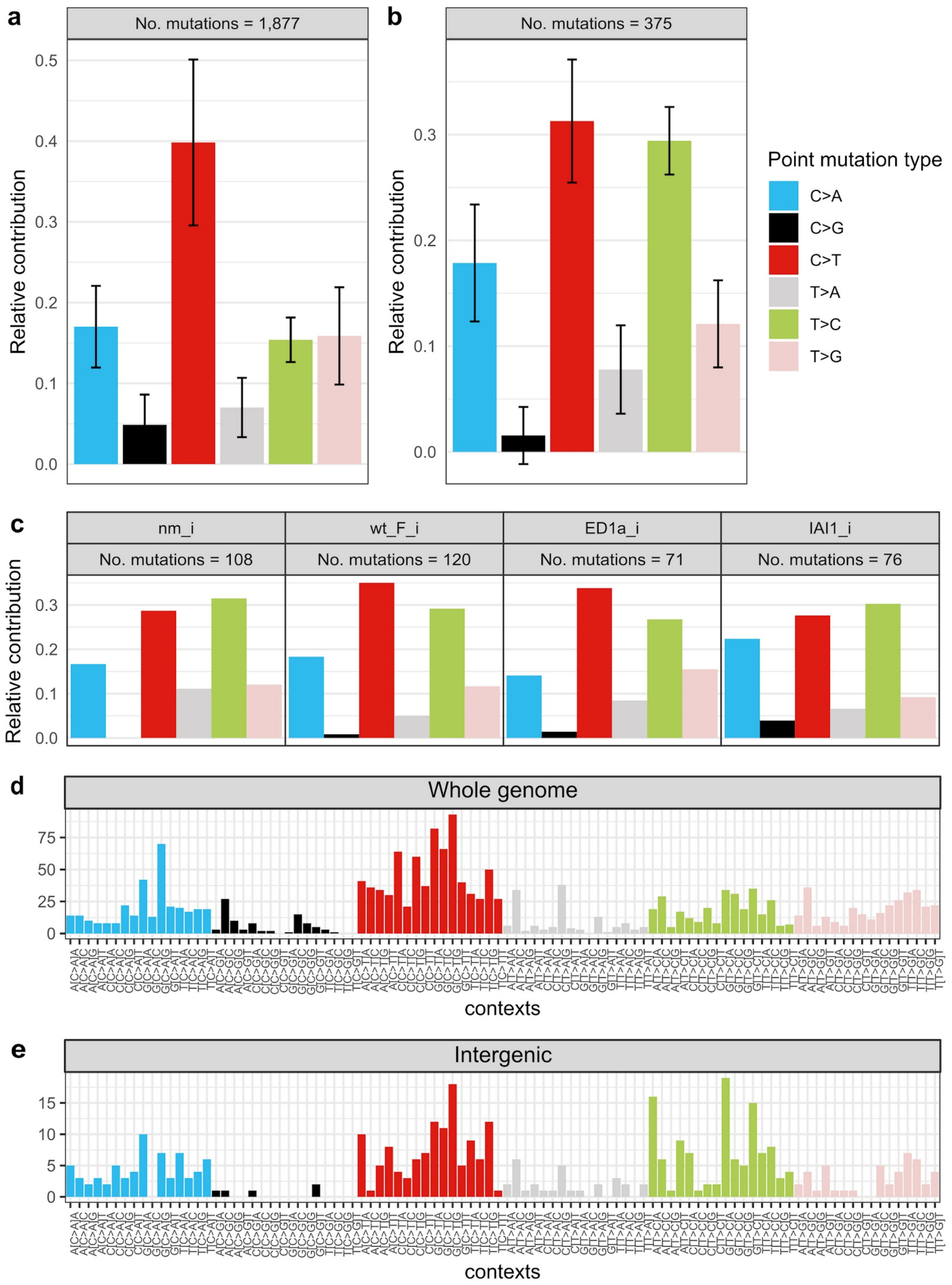


Supplementary Figure 5. Single nucleotide mutations in non-mutator laboratory strains. (a) Pooled SNPs in whole genome (b) Pooled SNPs in intergenic regions. (c) SNPs in intergenic regions by experiment. nm_i – non-mutator strains from LTEE, wt_F_i, ED1a_i, and IAI1_i – non-mutator strains from Foster et al., 2015. Substitutions with contexts in pooled data for the whole genome (d) and intergenic regions (e).

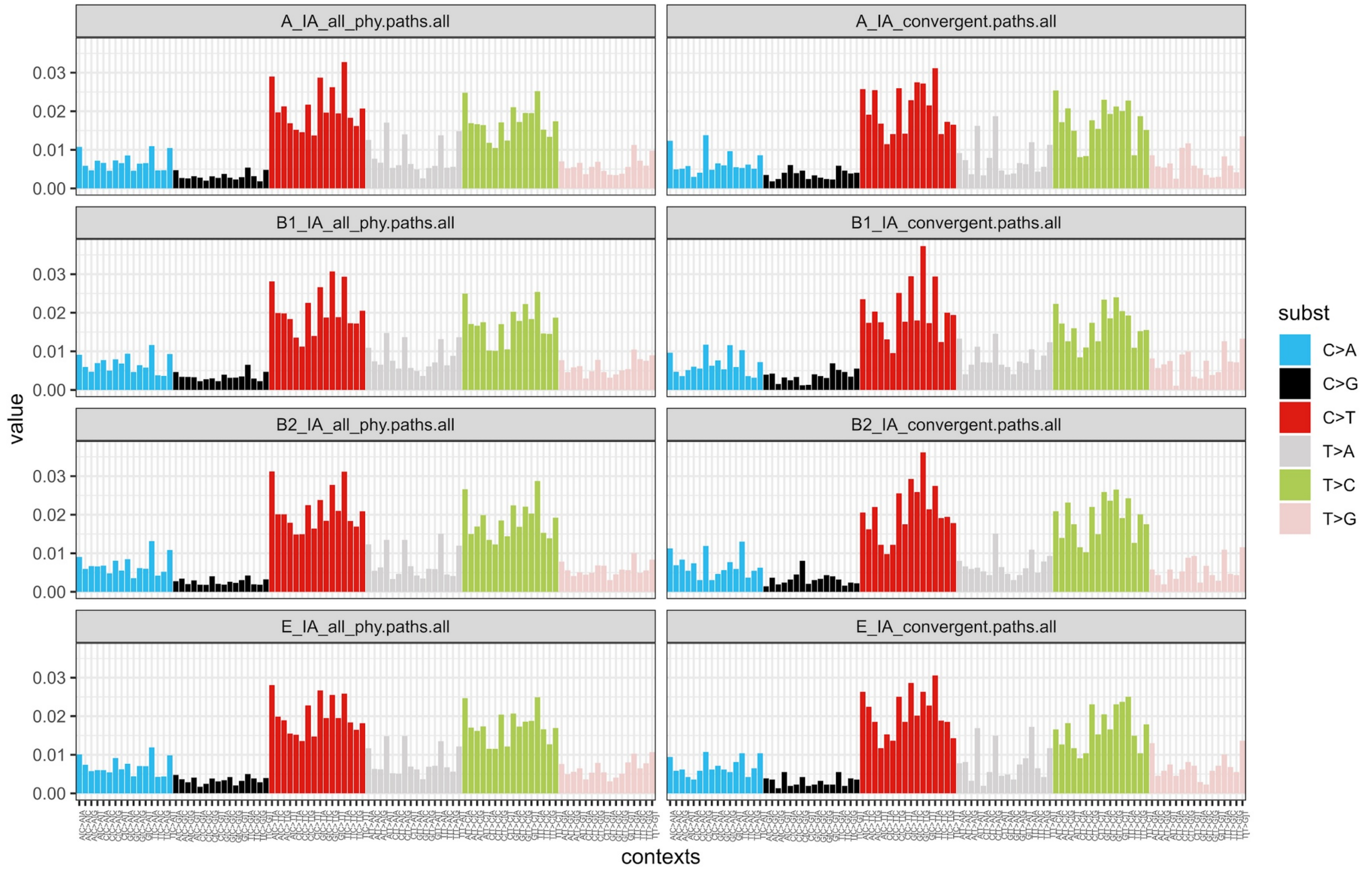


Supplementary Figure 6. Substitution profiles in divergence. Left column – all intergenic regions. Right column – all intergenic regions between convergent genes.

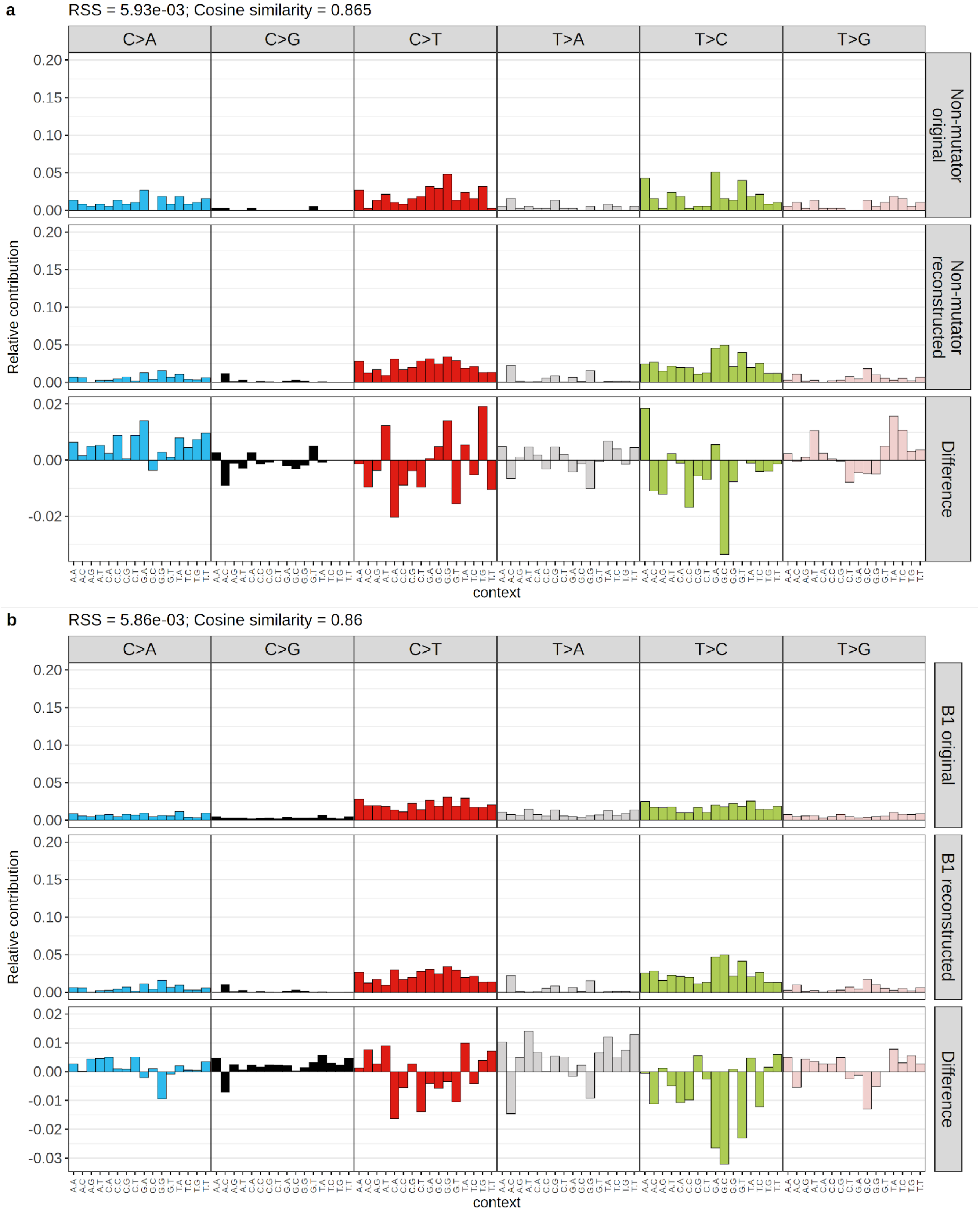


Supplementary Figure 7. Comparison between reconstructed and original mutational profiles for all intergenic regions in non-mutator strains (a) and natural lineages (b).

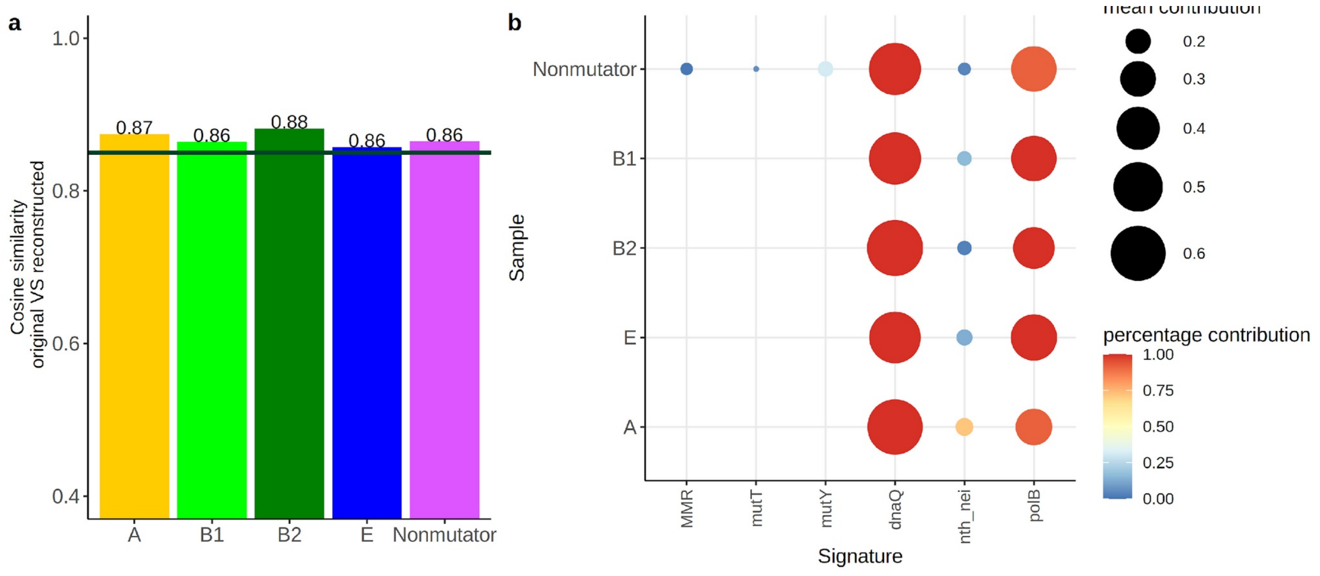


Supplementary Figure 8. Contribution of different mutational signatures to mutation accumulation in intergenic regions of convergent genes.

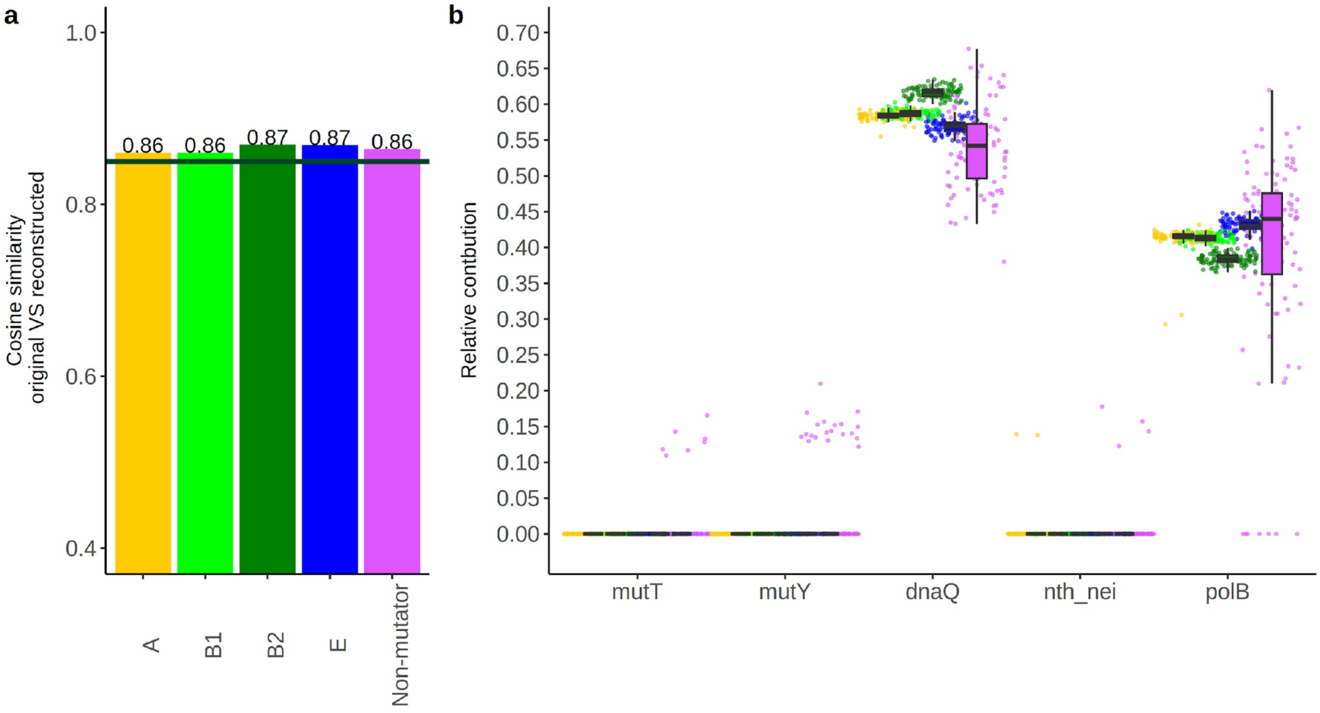


Supplementary Figure 9. Contribution of different mutational signatures to mutations accumulation in intergenic regions of all genes with recurrent mutations removed.
